# Single dose immunization with a COVID-19 DNA vaccine encoding a chimeric homodimeric protein targeting receptor binding domain (RBD) to antigen-presenting cells induces rapid, strong and long-lasting neutralizing IgG, Th1 dominated CD4^+^ T cells and strong CD8^+^ T cell responses in mice

**DOI:** 10.1101/2020.12.08.416875

**Authors:** Gunnstein Norheim, Elisabeth Stubsrud, Lise Madelene Skullerud, Branislava Stankovic, Stalin Chellappa, Louise Bjerkan, Katarzyna Kuczkowska, Elisabeth Müller, Monika Sekelja, Agnete B. Fredriksen

## Abstract

The pandemic caused by the SARS-CoV-2 virus in 2020 has led to a global public health emergency, and non-pharmaceutical interventions required to limit the viral spread are severely affecting health and economies across the world. A vaccine providing rapid and persistent protection across populations is urgently needed to prevent disease and transmission. We here describe the development of novel COVID-19 DNA plasmid vaccines encoding homodimers consisting of a targeting unit that binds chemokine receptors on antigen-presenting cells (human MIP-1α /LD78β), a dimerization unit (derived from the hinge and C_H_3 exons of human IgG3), and an antigenic unit (Spike or the receptor-binding domain (RBD) from SARS-CoV-2). The candidate encoding the longest RBD variant (VB2060) demonstrated high secretion of a functional protein and induced rapid and dose-dependent RBD IgG antibody responses that persisted up to at least 3 months after a single dose of the vaccine in mice. Neutralizing antibody (nAb) titers against the live virus were detected from day 7 after one dose. All tested dose regimens reached titers that were higher or comparable to those seen in sera from human convalescent COVID-19 patients from day 28. T cell responses were detected already at day 7, and were subsequently characterized to be multifunctional CD8^+^ and Th1 dominated CD4^+^ T cells. Responses remained at sustained high levels until at least 3 months after a single vaccination, being further strongly boosted by a second vaccination at day 89. These findings, together with the simplicity and scalability of plasmid DNA manufacturing, safety data on the vaccine platform in clinical trials, low cost of goods, data indicating potential long term storage at +2° to 8°C and simple administration, suggests the VB2060 candidate is a promising second generation candidate to prevent COVID-19.

## INTRODUCTION

In the period from December 2019 to December 2020, over 67 million cases and >1.5 million deaths due to COVID-19 disease have been reported (coronavirus.jhu.edu), and the non-pharmaceutical interventions required to limit the spread are having detrimental negative impacts on humanity. A safe COVID-19 vaccine able to prevent transmission of the SARS-CoV-2 virus and provide persistent protection against disease even in the elderly and immunocompromised is desirable, together with a product that can easily be administered, preferably as one dose regimen, and stored at +2-8°C or above (WHO 2020). This would require it to induce rapid and long-lasting neutralizing antibody levels and preferentially a Th1-biased CD4^+^ plus CD8^+^ T cell response.

The SARS-CoV-2 virus is a part of the Coronaviridae family, which apart from SARS-CoV and MERS-CoV mostly consists of human pathogens causing the common cold (Tse et al. 2020). Antibodies against the SARS-CoV-2 Spike (S) surface protein can block the virus from binding to the human cell receptor ACE2 and thus mediate virus neutralization (Barnes et al. 2020), and RBD is the primary target of S-specific neutralizing antibodies in convalescent sera (Robbiani et al. 2020). Neutralizing monoclonal antibodies isolated from convalescent COVID-19 patients have been shown to protect against SARS-CoV2 infection in challenge models in hamsters and non human primates (NHPs) (Baum et al. 2020), and several vaccines have induced nAbs that strongly correlated with reduction of viral load in NHPs (Klasse et al, 2020). This suggests nAbs to serve as a potential correlate of protection for COVID-19 disease, and most vaccine candidates in development focus on inducing neutralizing IgG antibodies against the S protein (Poland et al. 2020, Than Le et al. 2020). Studies on human COVID-19 patients also indicate a role of a balanced CD4^+^, CD8^+^ and neutralizing antibody responses in controlling the disease, with the CD8^+^ T cell responses likely required to avoid progression into severe COVID-19 disease in humans (Peng et al. 2020).

To address the need for a single-dose SARS-CoV-2 vaccine candidate ensuring persistent protection, on a platform enabling rapid adaptation to antigen changes, easily scalable manufacturing and a product that can be stable at +2-8°C, we developed a SARS-CoV-2 vaccine based on a DNA plasmid vaccine platform encoding a chimeric protein designed to enhance the antigen uptake through targeting the antigen to antigen-presenting cells (APC). The proteins are bivalent homodimers, each chain consisting of a targeting unit, a dimerization unit derived from the hinge and C_H_3 exons of human IgG3, and an antigenic unit (Fredriksen et al. 2006, Ruffini et al. 2010). The hinge region provides covalent binding of the two monomers via disulfide bonds while C_H_3 contributes to the dimerization through hydrophobic interactions. LD78β is an isoform of the human CC chemokine macrophage inflammatory protein-1α (MIP-1 α), and is suitable as a targeting unit due to its ability to attract APCs and deliver the antigen through chemokine receptors CCR1 and CCR5 (Ruffini et al. 2010). This leads to effective presentation of antigenic epitopes on MHC class I and MHC class II molecules to CD8^+^ and CD4^+^ T cells, respectively. This vaccine format has been shown to induce rapid, strong and dominant CD8^+^ cytotoxic T cell responses (Krauss et al. 2019, Ruffini et al. 2010).

We further built on the clinical experience of this vaccine format, where two similar LD78β chimeric vaccine products have been evaluated in clinical trials for the treatment of cancers. VB10.16 is a therapeutic HPV16-specific cancer vaccine which carries HPV16 E6 and HPV16 E7 in the antigenic unit and has been tested in a Phase 1/2a trial in patients with high grade cervical intraepithelial neoplasia (NCT02529930) and in an ongoing Phase 2 trial in patients with advanced or recurrent cervical cancer (NCT04405349). VB10.NEO is a fully personalized cancer neoantigen vaccine being tested in a Phase 1/2a trial in patients with multiple locally advanced or metastatic cancers (NCT03548467). Both vaccine candidates are delivered intramuscularly (i.m). using a needle-free jet injector (PharmaJet, U.S.). No significant safety concerns were detected to date and strong antigen specific immune responses were induced after vaccination (Hillemans et al. 2019, Krauss et al. 2019). Preclinical studies have shown that the vaccine platform can achieve both rapid (1 week) and long-lasting (at least 10 months) protection against influenza after a single dose (Grødeland et al. 2013, Lambert et al. 2016, Grødeland et al. 2019). The DNA plasmid vaccine platform exploited in these studies is a safe and highly versatile technology with intrinsic adjuvant effect designed for efficient delivery of antigen and inducing rapid, strong, broad and long-lasting immune responses. We here tailored the vaccine format against the SARS-CoV-2 virus (referred to as VB10.COV2) by including the Spike or RBD antigens into the antigenic unit of the LD78β chimeric vaccine construct, and evaluated anti-viral antibody and T cell responses in mice.

## RESULTS

### Design and characterization of VB10.COV2 proteins post transfection

Vaccine constructs were designed based on modifications of the antigens Spike and RBD. All synthesized DNA plasmids were evaluated for expression in human cells (HEK293), and subsequently evaluated for immunogenicity in BALB/c mice. VB10.COV2 DNA plasmids encoded either a short form of the SARS-CoV-2 RBD (“RBD short”, amino acids 331-524, i.e. 193 aa), a longer version (“RBD long”, amino acids 319-542, i.e. 223 aa), or the modified Spike protein (Figure 1a). These constructs were denoted VB2049 (RBD short), VB2060 (RBD long) and VB2065 (Spike), respectively (Figure 1b). The VB10.COV2 plasmids (Figure 1c) were transiently transfected into mammalian cells (HEK293), and the presence of the functional VB10.COV2 proteins in supernatant were measured by a sandwich ELISA using specific antibodies against the targeting, dimerization and antigenic units of the protein (i.e. LD78β, hIgG C_H_3 domain, the RBD domain or Spike protein). Reactivity confirmed successful expression and secretion, and a conformational integrity of all VB10.COV2 protein vaccine candidates (Figure 2). The expression level was found to be highest for VB2060 (RBD long), followed by VB2049 (RBD short) and VB2065 (Spike); indicating that VB2060 could also be secreted at higher levels from myocytes after i.m. vaccination.

**Figure 1.**
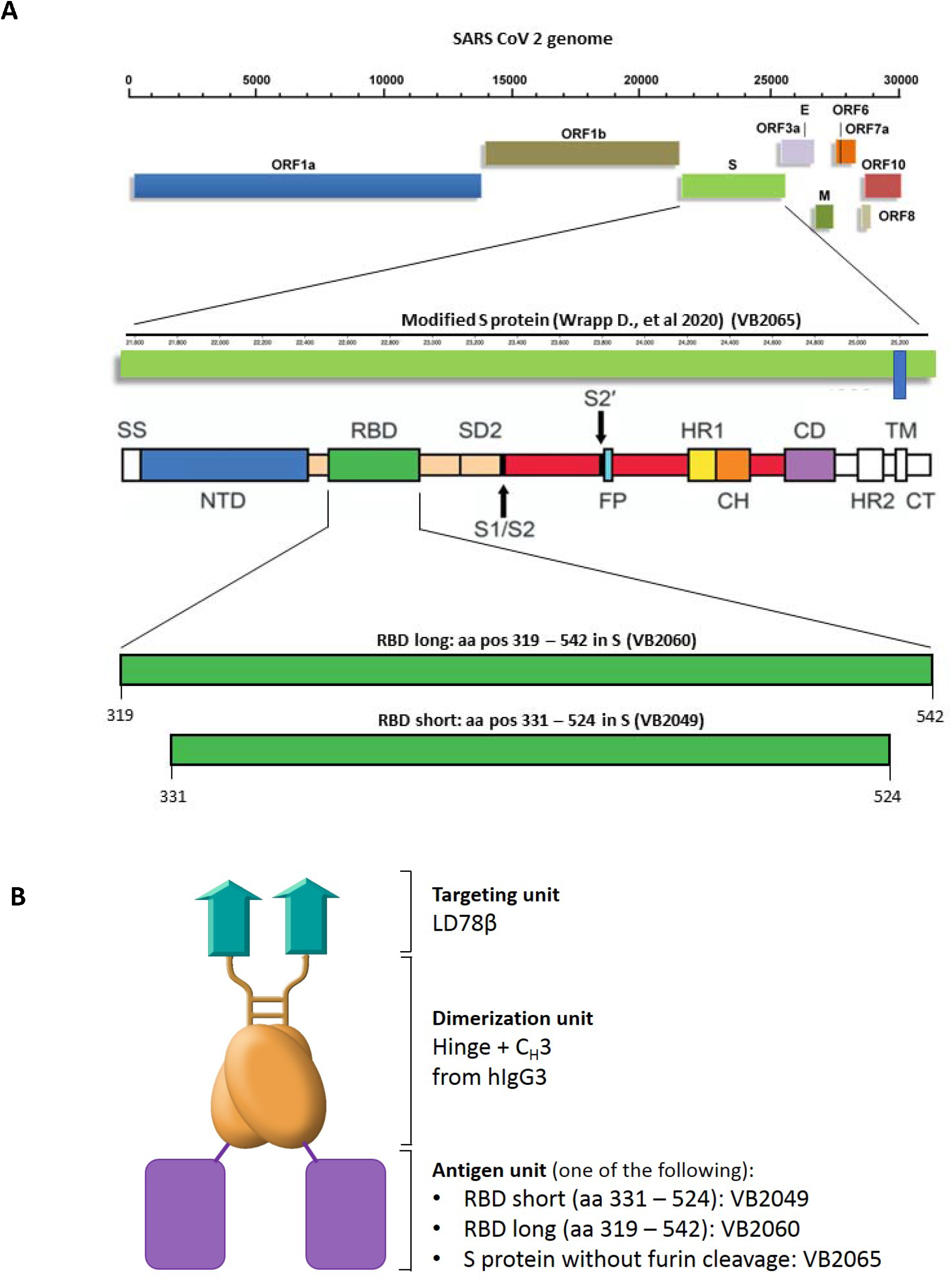
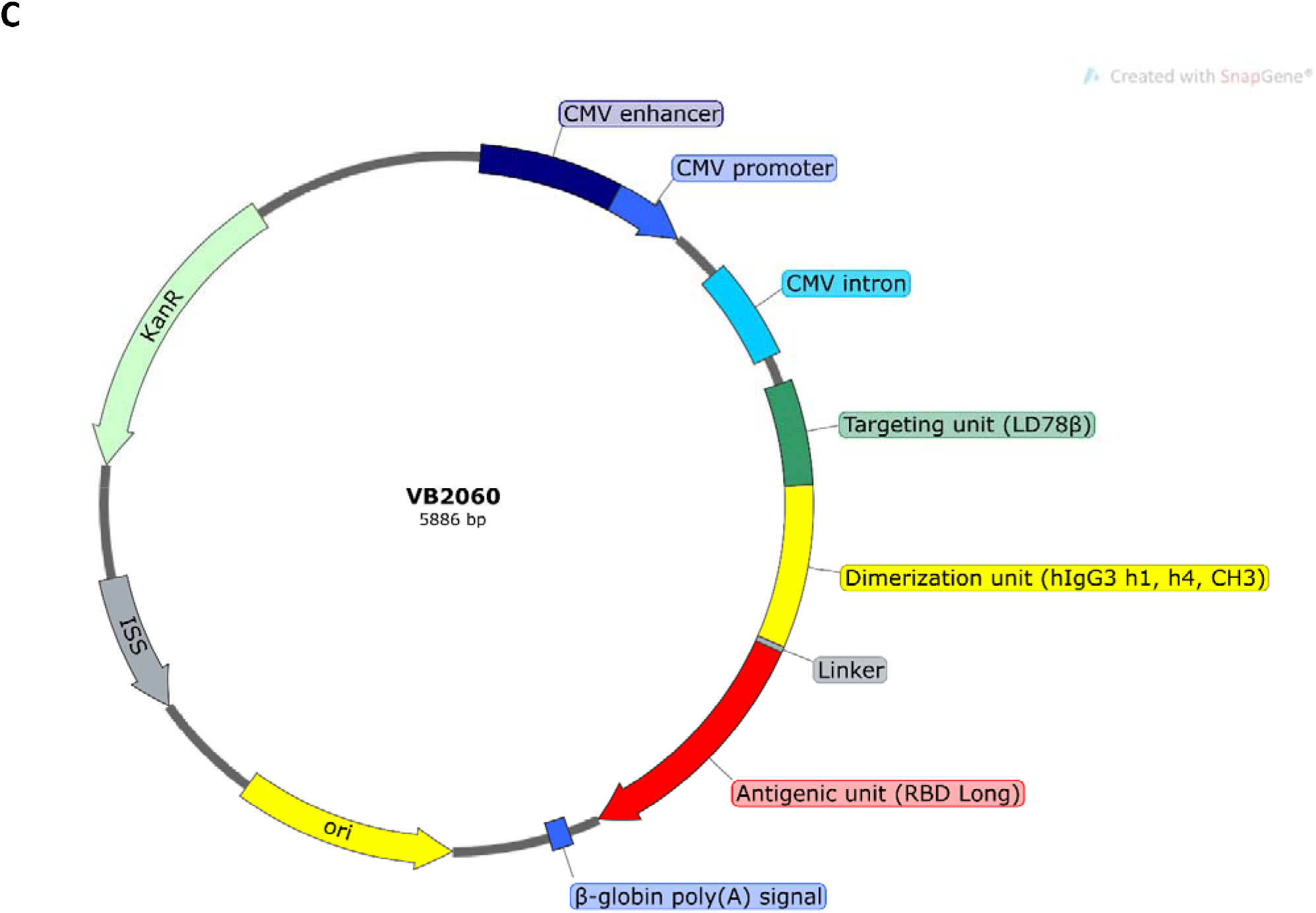
Alternative vaccine construct designs explored and characterized for immunogenicity in mice. **a** Schematic presentation of the SARS-CoV-2 genome, with the modified S protein (introduction of two prolines in the S2 subunit to stabilize S in prefusion conformation, and mutation of the furin cleavage site between S1 and S2 (Wrapp et al. 2020)) and amino acid positions of the “RBD long” and “RBD short” used in vaccine candidates indicated. **b** VB10.COV2 homodimeric proteins. Each chain of the dimer contains a N-terminal LD78β targeting unit (turquoise), a dimerization unit (yellow) composed of a shortened IgG hinge and C_H_3 domain from human γ3 chains, and a C-terminal antigen unit genetically linked to the dimerization unit. LD78 is an isoform of the human CC chemokine macrophage inflammatory protein-1α (MIP-1 α). Antigens encoded by the different constructs: VB2065; codon-optimized stabilized S protein without furin cleavage. VB2060; RBD (long, aa 319 – 542). VB2049; RBD (short, aa 331 – 524). **c** Structure of plasmid construct encoding VB2060, based on the expression vector pUMVC4a and with the homodimeric RBD protein insert shown.

**Figure 2.**
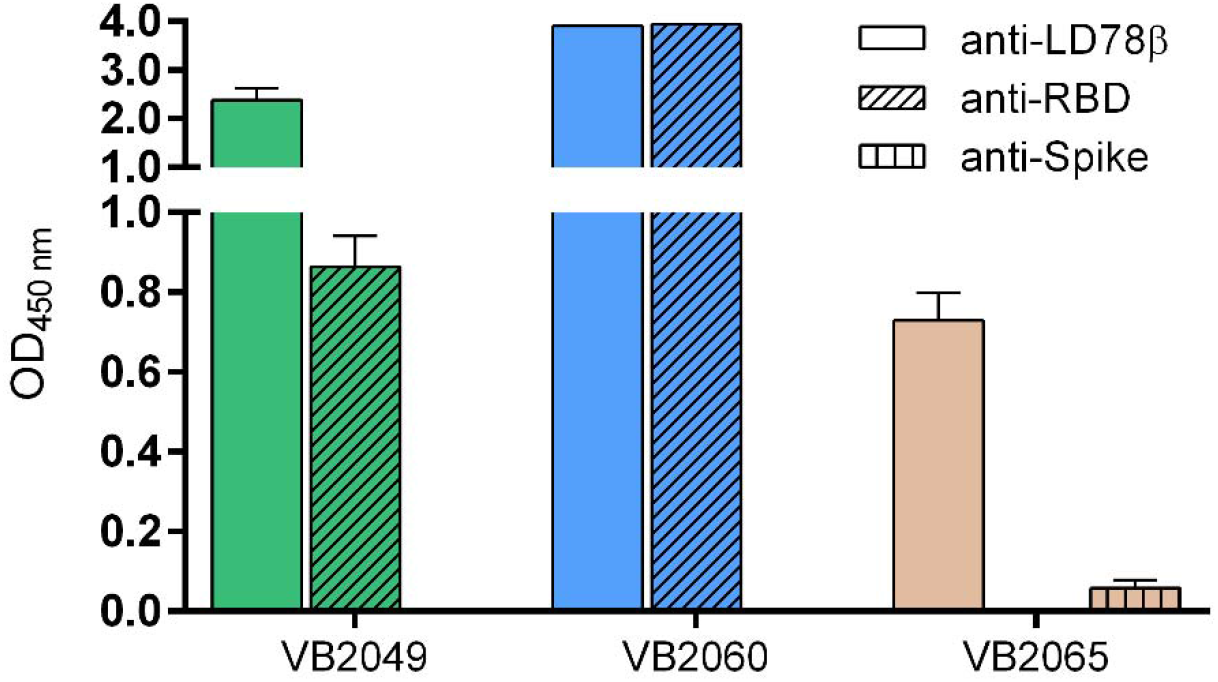
VB10.COV2 construct proteins (VB2049, VB2060 and VB2065) produced and secreted as functional homodimers 3 days after transfection of HEK293 cells. Conformational integrity of the epitopes expressed in the constructs was confirmed by binding to antibodies detecting the human LD78β, human IgG C_H_3 domain, the RBD domain or Spike protein in ELISA.

### Humoral immune responses to SARS-CoV-2 antigens induced in mice

The constructs were compared for the ability to induce anti-RBD IgG. All three candidates, when administered at a dose of 50 μg, evoked strong IgG responses against RBD in mice. The antibodies induced by the construct encoding RBD-based antigens (VB2049 and VB2060) were detected already at day 7 (Figure 3), while for candidates encoding S protein (VB2065) antibodies were first detected at day 14. VB2060 seemed to induce higher, and more rapid anti-RBD antibody responses compared to VB2049 and VB2065 (Figure 3). The antibody response induced by VB2060 was further characterized, and demonstrated anti-RBD IgG as early as day 7 post single vaccination; even at a low dose (2.5 μg) (Figure 4a), as well as a consistent dose-response (Figure 4b)The antibody levels peaked at day 28 (10^5^ endpoint titer) after a single dose and achieved high levels for at least 89 days (Figure 4a). For VB2060, the peak and durability of the response were further increased (>10^6^ endpoint titer) following a two dose regimen (days 0 and 21, at both 50 μg and 25 μg) compared to the single dose group. Limited added benefit was observed at day 99 in mice that received a boost vaccination at day 89 (Figure 4a). A second experiment confirmed a clear tendency of a dose-dependent response in the range of 3, 6, 12.5 and 25 μg of VB2060 (Figure 4b), in particular on day 7, with high Ab levels reached for all groups from day 14 until day 28, both with a single and two-dose regime. The responses were significantly different between the groups of mice vaccinated with 3 μg vs. 12 μg, 3 μg vs. 25 μg, and 6 μg vs. 25 μg.We further tested the kinetics of RBD-specific IgG in bronchoalveolar lavage (BAL) from mice vaccinated once or twice with different doses VB2060 (Figure 4c). RBD-specific IgG was found in BAL at the earliest time point tested (day 14) even with the lowest dose, and the levels increased with dose and over time (Figure 4c).

**Figure 3.**
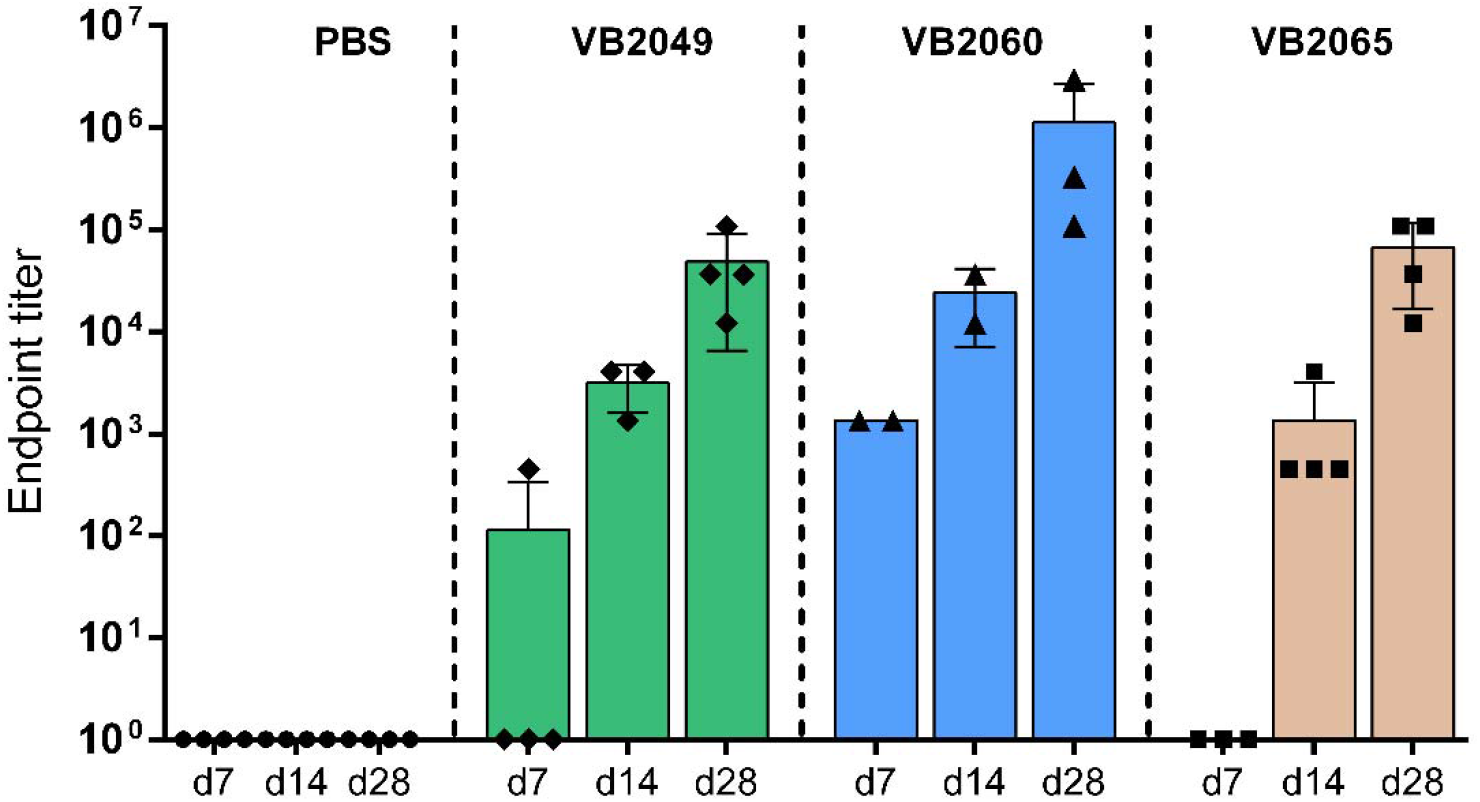
Anti-RBD IgG responses in mice vaccinated with 50 μg of VB10.COV2 candidates (VB2049, VB2060 or VB2065). Mice were vaccinated by i.m. administration of DNA immediately followed by electroporation of the injection site at day 0 and day 21. Sera obtained at day 7, 14 and 28 post first vaccination with VB2049, VB2060 or VB2065 were tested for anti-RBD IgG antibodies binding the RBD protein. Data are shown as mean ± SEM with individual values (*n* = 2-4).

**Figure 4.**
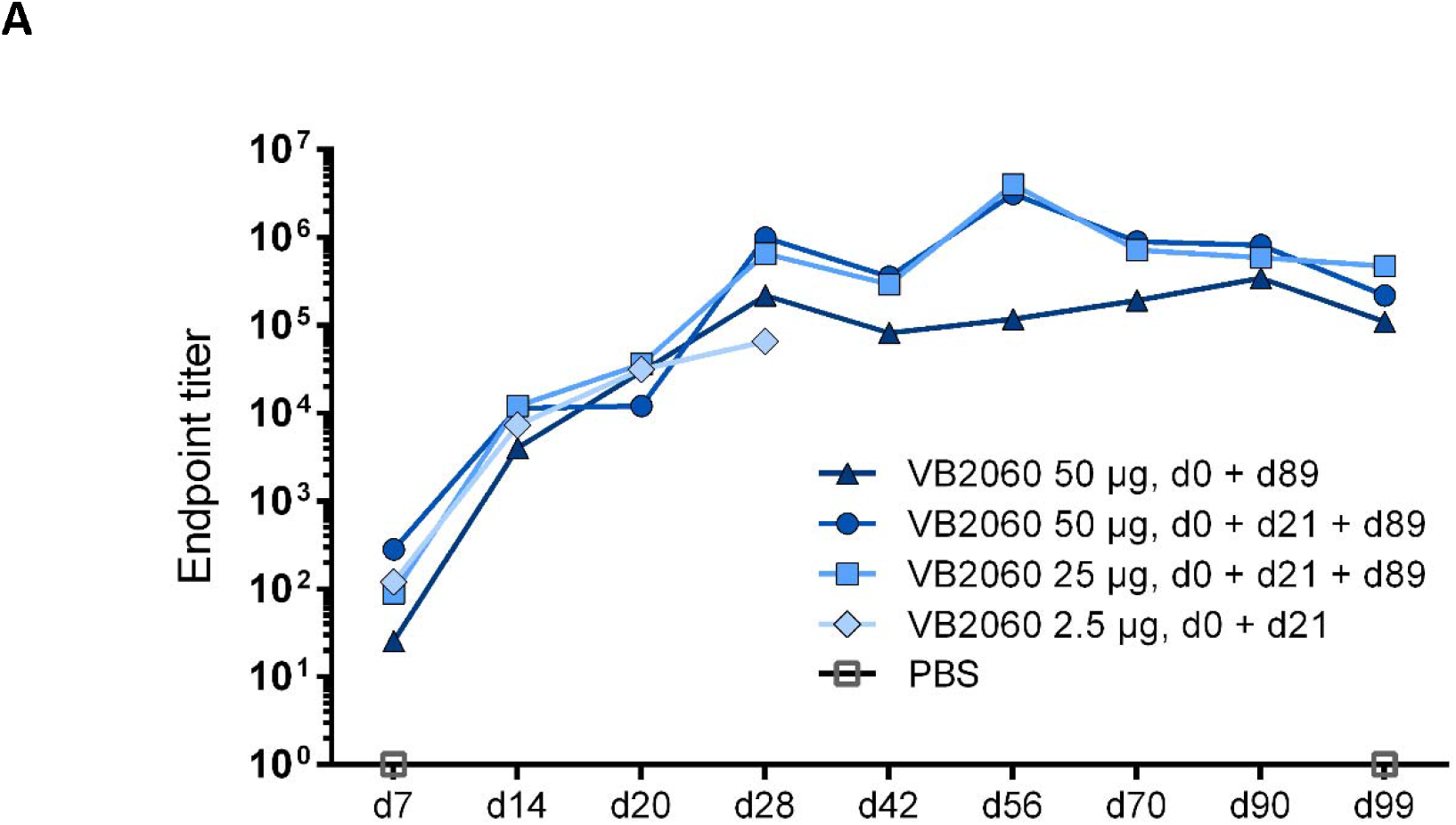
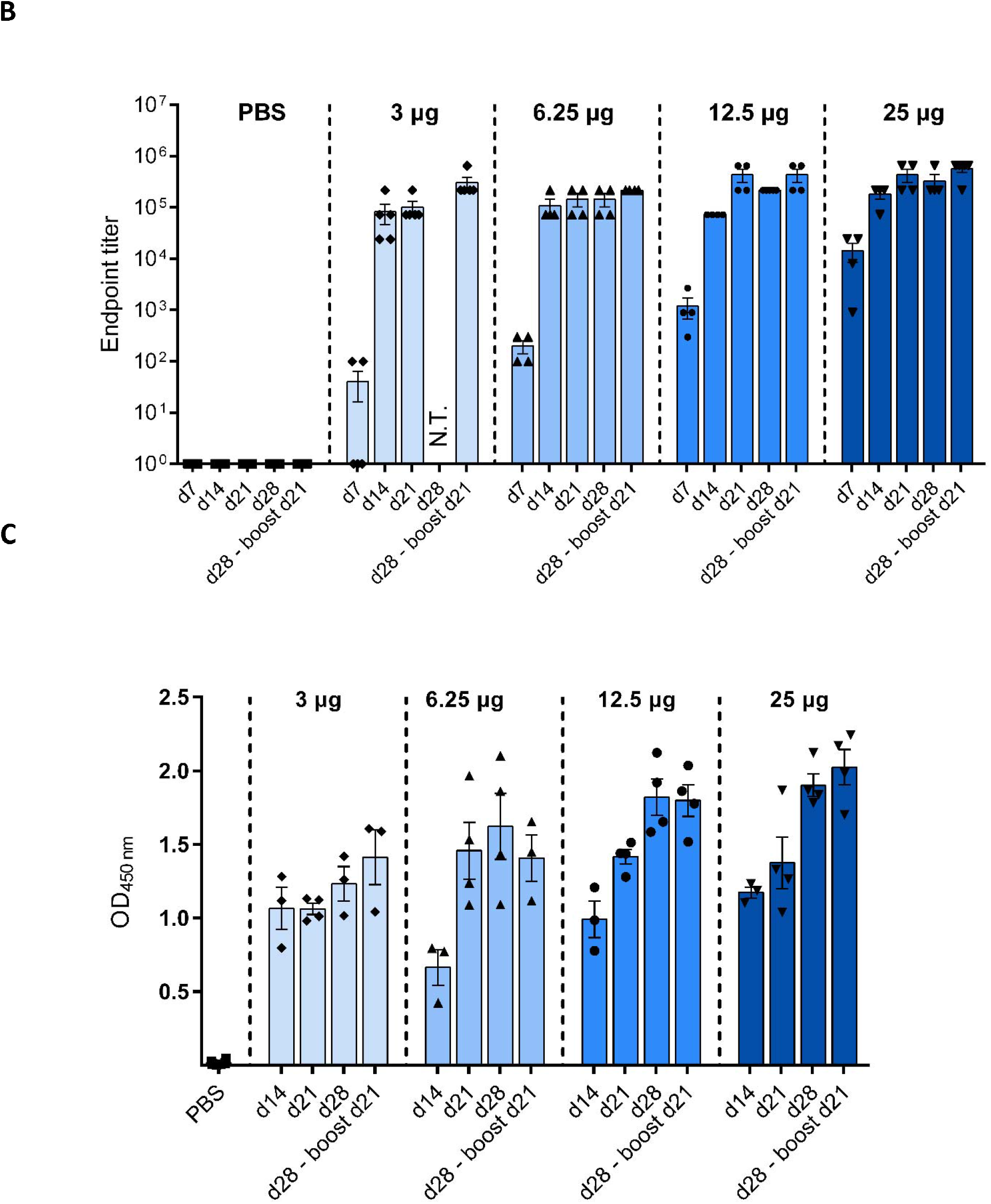
Kinetics of anti-RBD IgG response in mice vaccinated with VB10.COV2 VB2060. Mice were vaccinated by i.m. administration of VB2060 immediately followed by electroporation of the injection site. Dose time points and dose levels are indicated. **a** Serum anti-RBD IgG assessed until day 99 post first vaccination. The results are shown as a mean of two independent ELISA experiment. Group size is *n* = 5 until day 70, *n* = 2-3 at the two last time points (days 90 and 99). **b** Serum anti-RBD IgG responses in mice vaccinated with 1 (day 0) or 2 doses (days 0 and 21) of either 3, 6, 12.5 or 25 μg of VB2060, measured weekly for up to 4 weeks (*n* = 4-5 mice per group, data shown as mean ± SEM with individual values, NT; not tested. **c** Anti-RBD IgG responses in bronchoalveolar lavage (BAL) from mice immunized with 1 (day 0) or 2 doses (days 0 and 21) of either 3, 6, 12.5 or 25 μg of VB2060, measured at 14 and 21 and 28 days after first vaccination and 7 days post boost. Data are shown as mean ± SEM with individual values (*n* = 3-4).

Sera were also assessed in a live SARS-CoV-2 virus neutralization assayOne dose of 50 μg of VB2060 induced strong and long lasting neutralizing antibodies, and was sufficient to induce detectable neutralizing activity already at day 7 which peaked at day 28 with no signs of decline at day 99 (ND_50_ 10^4^) (Figure 5a). Strong neutralizing antibody responses were seen in pools from vaccinated mice with all three candidates VB2049, VB2060 and VB2065 (Figure 5b, Supplemental figure S1). Serum pools from groups mice subjected to various doses and dosing regimens were tested for nAbs at selected time points (day 28, 90 and 99) for VB2060 and VB2049 (Figure 5b). Two doses of 2.5 μg of VB2060 resulted in titers >10^3^ at day 28 (data not shown). All regimens reached higher or comparable titers to the NIBSC convalescent plasma reference serum 20/130 from day 28 and until the end of the experiment. Independent of the dose, the strongest response was observed at day 99 (after boost at day 89), showing induction of long-lasting, neutralizing antibody responses with VB2060. Taken together, VB2060 was found to be superior to VB2065 and VB2049 in inducing rapid and high levels of neutralizing antibodies.

**Figure 5.**
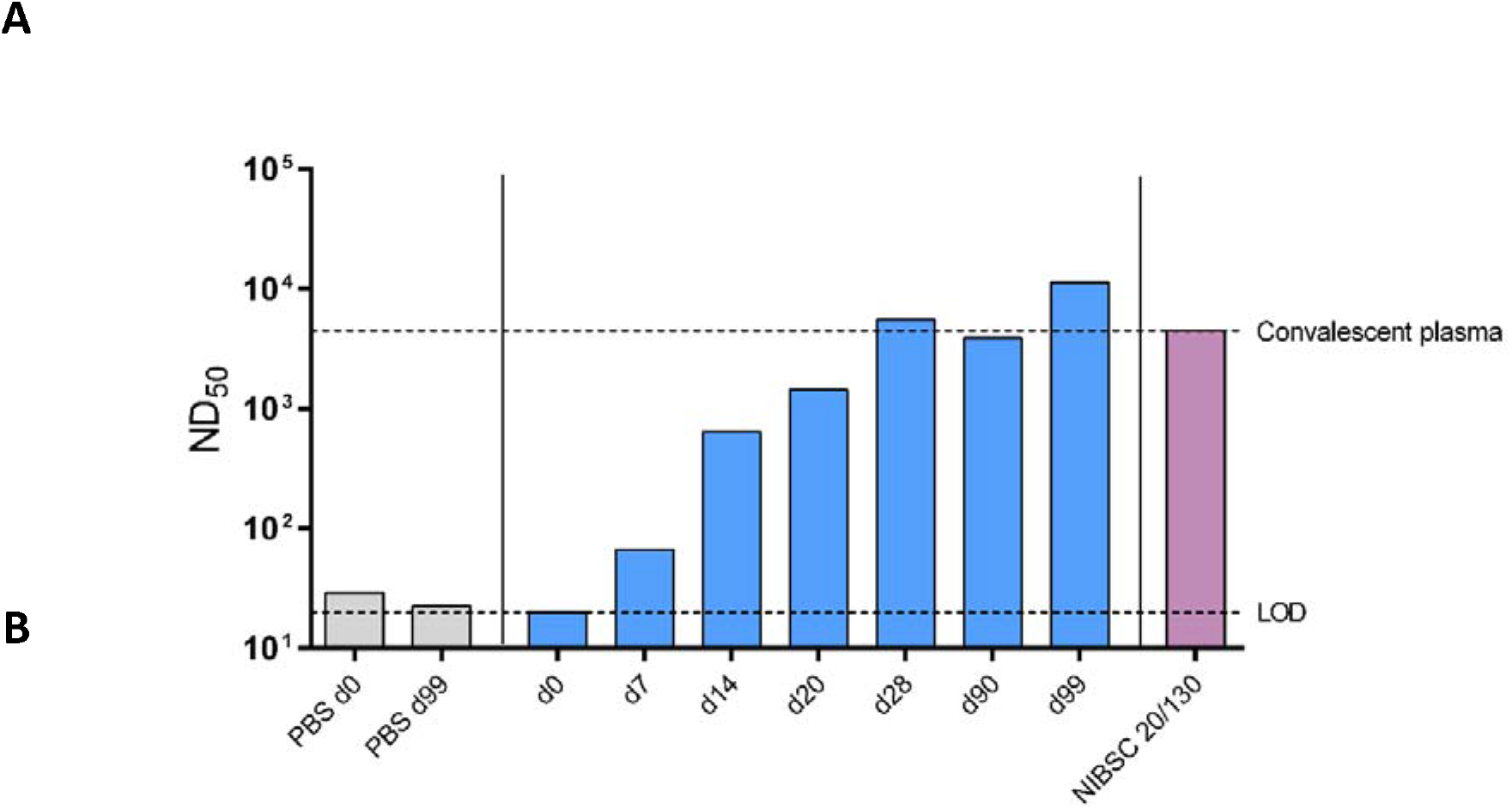
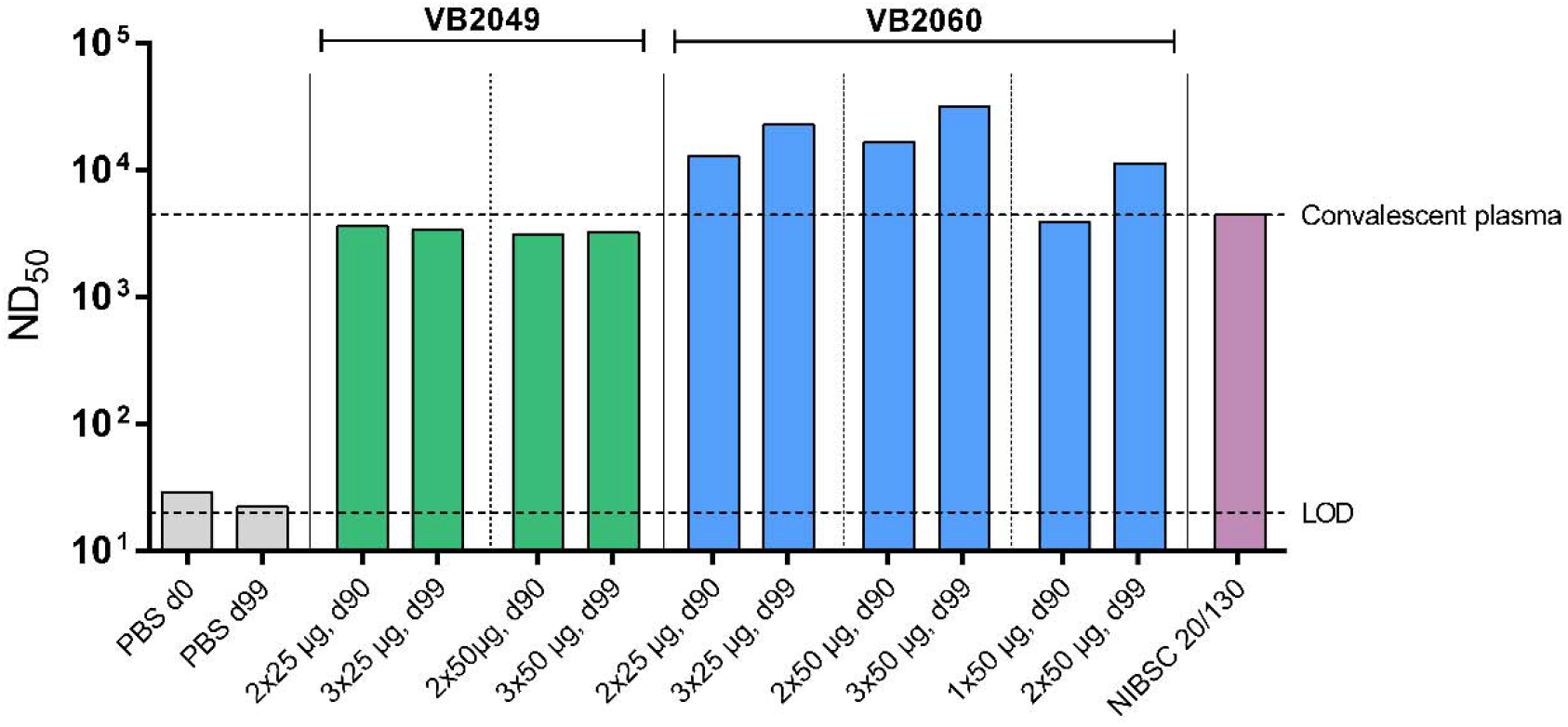
Neutralizing antibody responses in sera after immunization with VB2060. Bars represent endpoint titers for pooled sera assessed in live virus neutralization against homotypic SARS-CoV-2 live virus strain Australia/VIC01/2020. Lower dashed line indicate the limit of detection (LOD) of the assay, and the upper dashed line the titer value for the convalescent serum reference NIBSC 20/130 (ND_50_ endpoint titer 4443). **a** Neutralizing antibody responses in sera after immunization with one 50 μg dose of VB2060. **b** Neutralizing antibody responses in sera from mice immunized with 25 μg or 50 μg of VB2060 or VB2049 vaccines either as a one dose regimen (day 0) or two dose regimen (days 0 and 21) and boosted at day 89.

### Evaluation of the magnitude and specificity of T cell responses after vaccination

Overall, the VB10.COV2 constructs all induced strong, dose-dependent T cell responses after vaccination, that increased over time. The responses were dominated by CD8^+^ T cells and accompanied by significant, but weaker CD4+ T cell responses. Vaccination with 25 μg of VB2060 induced T cell responses as early as day 7 (~550 per 10^6^ splenocytes), (Figure 6a). The T cell responses increased until day 28 (Figure 6b) and were still found to persist for at least 90 days after vaccination with 50 μg of VB2060 (~5000 SFU/10^6^ splenocytes), with a strong boost effect at day 99; 10 days after a new booster dose was administered at day 89 (~20 000 SFU/10^6^ splenocytes) (Figure 6c). We further sought to characterize the epitopes recognized by the T cells by stimulating with individual 15-mers overlapping with 12 amino acids in splenocytes depleted for either CD4 or CD8 T cell populations. Strong (up to ~4000 SFU/10^6^ cells) CD8^+^-T cell responses against 9 peptides were observed. RBD-specific CD4^+^ responses were also detected against 7 peptides, but of a lower magnitude (up to ~1000 SFU/10^6^ cells) and fewer epitopes (Figure 7). The amino acid sequence of the overlapping peptides indicated a reactivity against 4 distinct MHC class I-restricted epitopes and 3 MHC class II-restricted epitopes (Supplemental figure S2 and table S1) in RBD.

**Figure 6.**
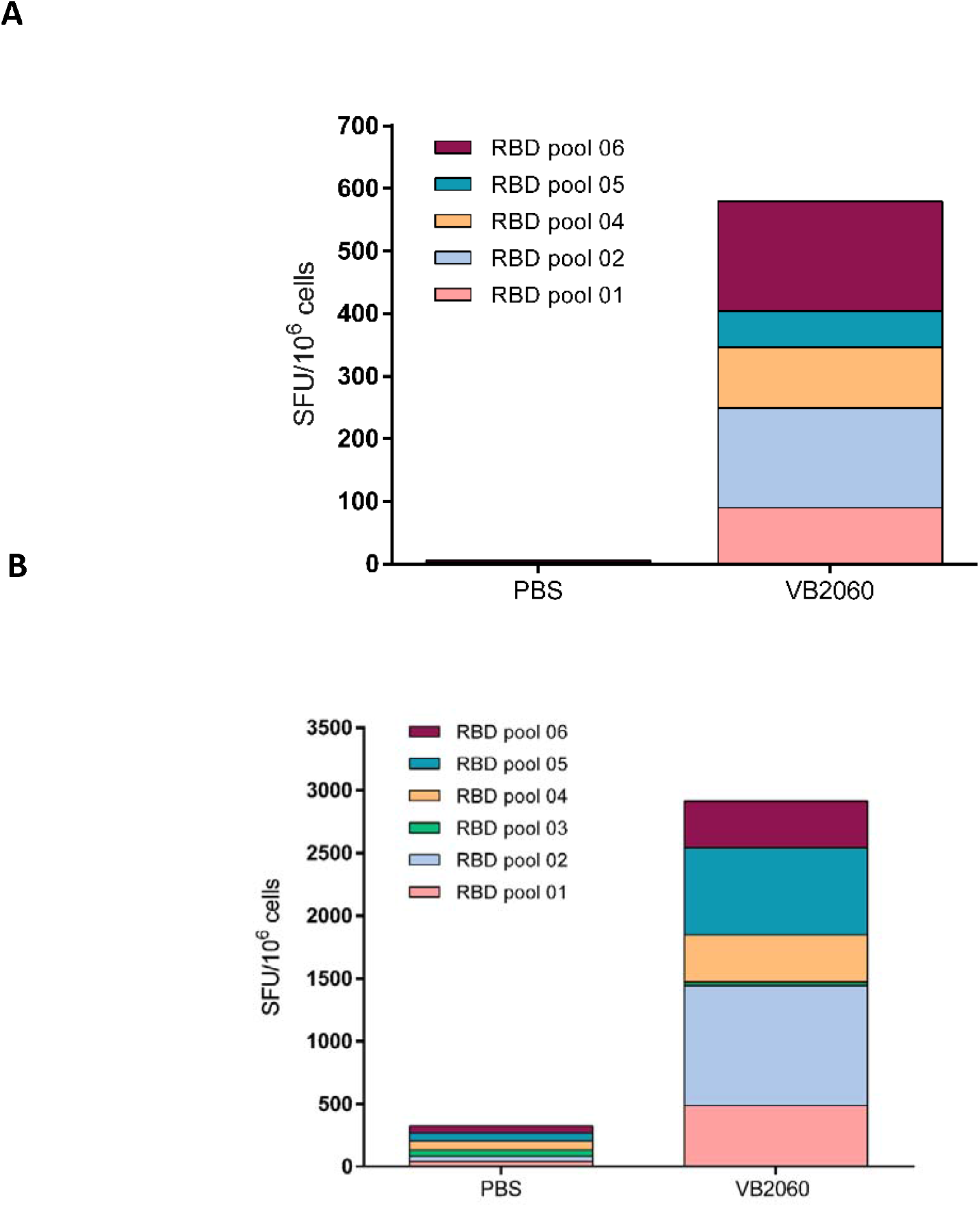
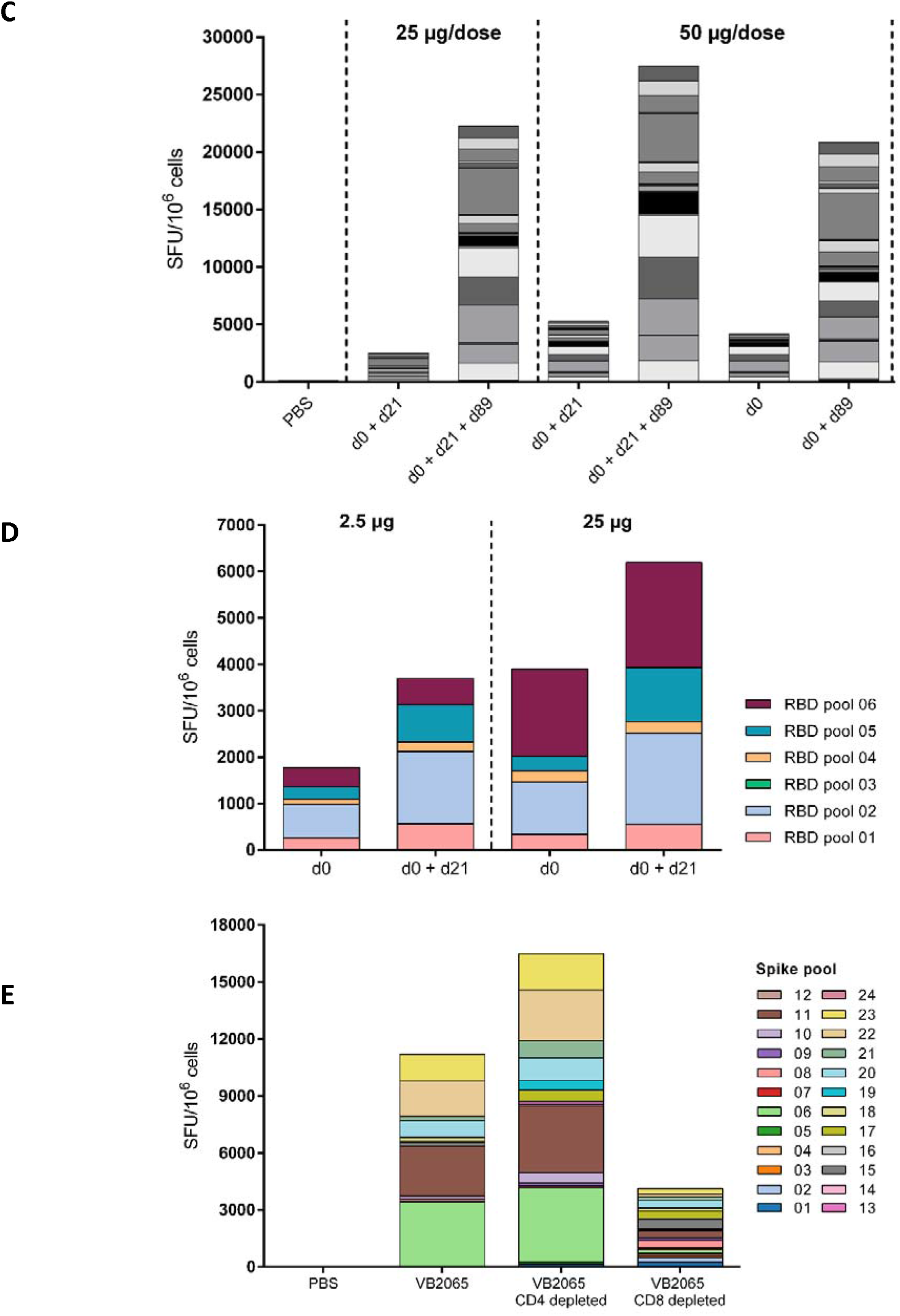
T cell responses induced with different doses and number of doses of VB10.COV2 DNA vaccines VB2049, VB2060 or VB2065. Splenocytes were harvested at day 7, day 14, day 28, day 90 or day 99 post first immunization and/or at day 7 or day 10 post boost vaccination at day 21 or day 89, respectively, and the total number of IFN-γ positive spots/1×10^6^ splenocytes after restimulation with overlapping RBD or Spike peptide pools or individual RBD peptides were determined by IFN-γ ELISpot. **a** Mice (*n* = 5) were i.m. vaccinated once with 25 μg VB2060 plasmid and spleens harvested at day 7. **b** Mice (*n* = 5) were i.m. vaccinated at day 0 and day 21 with 2.5 μg VB2060 DNA plasmid and spleens harvested at day 28. **c** Persistence of RBD-specific T cell responses after vaccination with VB2060 plasmid, measured by IFN-γ + ELISpot assay tested for 61 individual RBD peptides. Responses at different dose levels (25 μg or 50 μg); for the 25 μg dose level, responses were measured at day 90 after a two dose regimen (days 0 and 21) and 10 days post boost (i.e. day 99). For the 50 μg dose level, responses were measured at day 90 after a two dose regimen (days 0 and 21) and 10 days post boost (i.e. day 99), as well as at day 90 after a one dose regimen (day 0) and 10 days post boost (i.e. day 99). **d** Mice (*n* = 5) were i.m. vaccinated at day 0 and day 21 with 2.5 μg or 25 μg VB2049 DNA plasmid and spleens harvested at day 14 (for mice vaccinated at day 0) or at day 28 (for mice vaccinated at day 0 and day 21). **e** Mice (*n* = 8) were i.m. vaccinated at day 0 and day 21 with 50 μg VB2065 DNA plasmid and spleens harvested at day 28. CD4^+^ and CD8^+^ depletion of splenocytes was performed to elucidate the epitope specific distribution of responses among CD4^+^ or CD8^+^ T cell populations.

**Figure 7.**
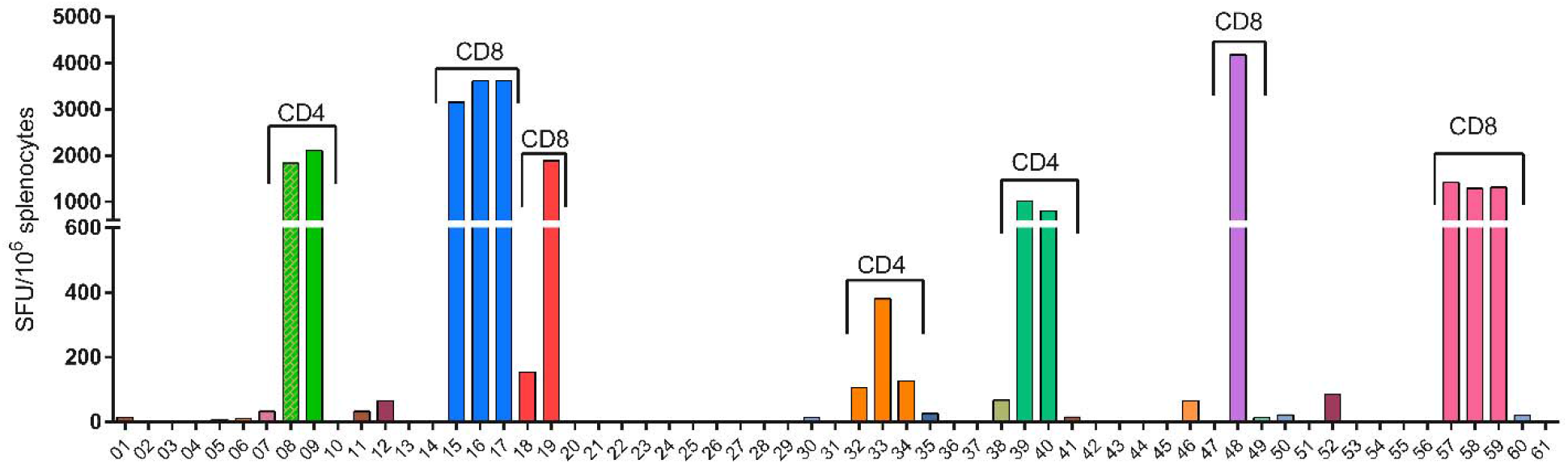
Identification of CD4^+^ and CD8^+^ RBD-specific T cell epitopes at day 99 post first vaccinations of VB2060 (*n* = 2). Splenocytes were harvested at day 99 from mice vaccinated at day 0, 21 and 89 and stimulated for 24 hours with 61 individual RBD peptides (15-mer peptides overlapping by 12 aa from SARS-CoV2 RBD domain) and the number of IFN-γ positive spots/1×10^6^ splenocytes were detected in an ELISpot assay. Indications of CD4^+^ and CD8^+^ specific responses are derived from an CD4 and CD8 depletion experiment with VB2049 with the same 61 peptides (Supplemental Figure S2).

Strong T cell responses against the RBD domain of SARS-CoV-2 were detected in spleens from mice vaccinated with one or two doses of both 2.5 μg or 25 μg VB2049 (Figure 6d). Depending on dose level and the number of doses, the response ranged from ~1800 to 6000 SFU per 10^6^ cells in splenocytes sampled 2 weeks after 1^st^ dose or 1st week post-boost-vaccination at day 21 and stimulated separately with 6 peptide pools spanning RBD. The response was strong already at 14 days post 1^st^ vaccination even with a low dose (2.5 μg DNA) and was boosted by day 28 in groups receiving a 2^nd^ vaccination at day 21 (in a dose-dependent manner, Figure 6d). When comparing T cell responses induced by two doses of 2.5 μg of either VB2060 or VB2049, VB2049 induced stronger responses than VB2060 (~3800 versus ~2600 SFU/10^6^ cells) (Supplemental figure S3). As predicted, VB2065 induced a broader, stronger total T cell response than VB2049 and VB2060 due to the larger antigen with strong, CD8+ dominating T cell responses, accompanied by broad, weaker CD4+ responses. (Figure 6e, Supplemental table S2).

### Evaluation of polyfunctional T cell responses and Th1/2/17 cytokine profile in splenocytes

Splenocytes from mice vaccinated with 2 doses of 2.5μg VB2060 were harvested on day 28 and restimulated with RBD peptide pools and the cell culture supernatants were analyzed for Th1 (IFN-γ, TNF-α, IL-12), Th2 (IL-4, IL-5) cytokines and IL-6 (Figure 8). The response was dominated by IFN-γ and TNF-α, and minor quantities of IL-4, IL-5, IL-6 or IL-12 p70 were detected (Figure 8). This indicates that T cell responses showed strong Th1 bias when characterized one month after immunization, while the Th2 responses were minimal.

**Figure 8.**
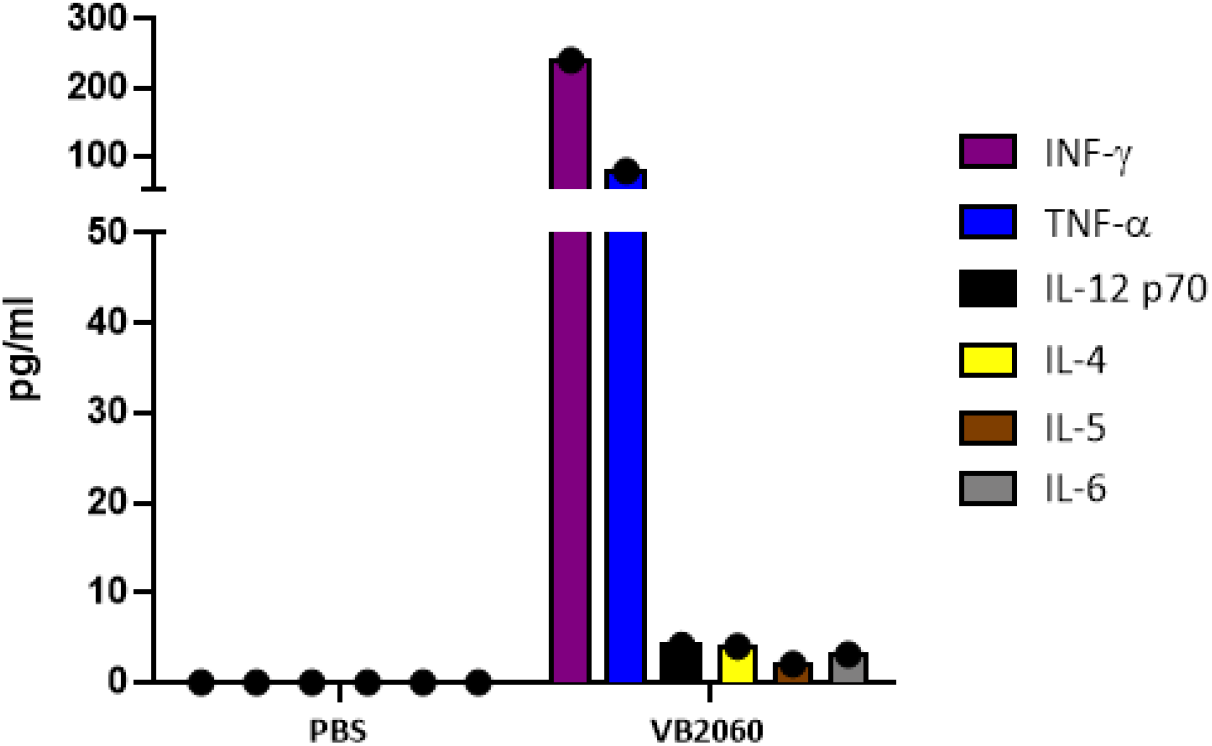
Characterization of the Th1 (IFN-γ, TNF-α, IL-12p70) and Th2 (IL-4, IL-5) cytokines in cell culture supernatant. Splenocytes from mice vaccinated with 2.5 μg VB2060 at day 0 and day 21 were restimulated for 16 h with RBD peptide pools (1-6) on day 28. Cytokine concentrations were measured using bead based immunoassay (ProcartaPlex).

In depth, T cell analysis of splenocytes from mice vaccinated with 1 or 2 doses of VB2060 were performed using flow cytometry on day 90 post prime vaccination. In VB2060 vaccinated mice, RBD specific T cell responses were dose dependent and varied between 2.07% to 6.3% of CD8^+^ T cells, and 1.08% to 3.3% of CD4^+^ T cells (Figure 9). CD8^+^ T cell responses were dominated by single or combined production of IFN-γ and TNF-α indicating an effective RBD-specific cytotoxic response (Figure 9B).

**Fig. 9.**
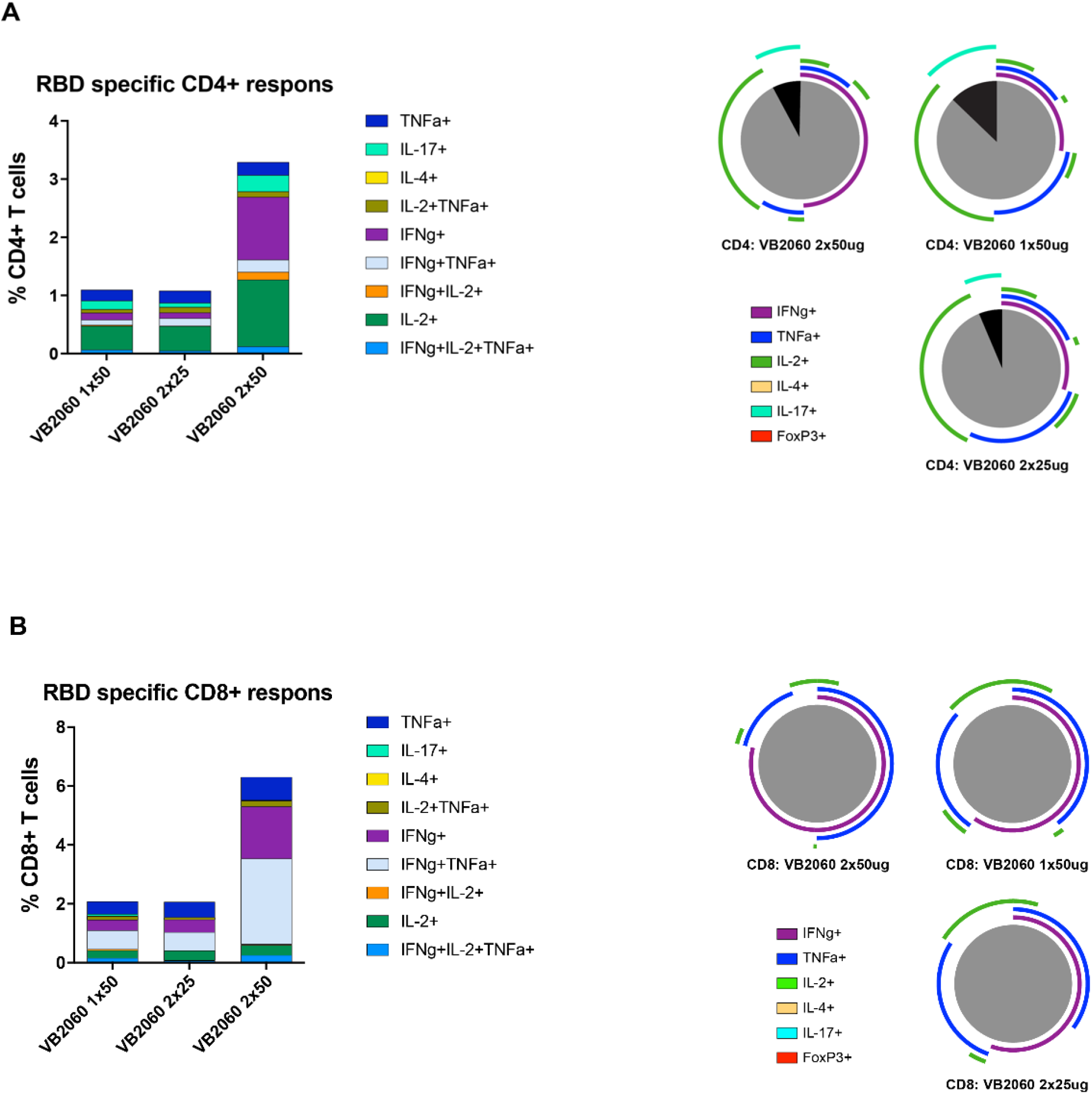
RBD specific multifunctional T cell responses in mice vaccinated with VB2060. Mice were vaccinated on day 0 and 21 and T cell responses were analyzed on day 90 using multiparameter flow cytometry. **a** Percent of CD4^+^ and **b** CD8^+^ T cells responding to RBD stimulation. Percent of RBD specific cells is shown in bar graphs. Cells expressing one marker, or combinations of multiple markers are shown as percent of parent population. Pie charts show cytokine profile. CD4+ T cell responses presented with Th1 (grey) / Th17 (black) bias. Pie charts were made using Spice software.

The CD4^+^ RBD-specific T cells displayed a typical Th1 profile indicated by production of IFN-γ, TNF-α, IL-2 or a combination of these (Figure 9A). The RBD specific multifunctional CD4^+^ T cells (expressing 3 cytokines simultaneously) also showed a dose dependent response, where the highest response was observed in 2 × 50 μg VB2060 vaccinated mice (0.13%) 90 days after the initial vaccination. Presence of other markers like IL-4 (Th2 polarization), IL-17 (Th17) and FoxP3 (Treg) was also tested. A minor population of CD4^+^ cells expressed the Th17 cytokine, IL-17 (Figure 9A), while no significant IL-4 or FoxP3 were detected, thus revealing a combination of Th1 and Th17 responses with strong bias towards Th1. Taken together, these data showed that the VB2060 vaccine induced strong Th1 responses which persisted up to at least 3 months (90 days).

## DISCUSSION

We here demonstrate that when encoding SARS-Cov-2 RBD in a chimeric DNA vaccine construct, the expressed chimeric fusion protein that binds to chemokine receptors on APC, presents RBD in a conformation able to induce rapid RBD-specific and neutralizing antibody responses. IgG against RBD was shown to correlate with neutralizing antibody activity in humans (Poland et al. 2020), and in non-human primates neutralizing antibodies against Spike was shown to function as a correlate of protection (CoP) (Chandrashekar et al. 2020). The anti-RBD IgG detected in lungs after vaccination with VB2060 may contribute to the local virus neutralization serving the first line of protection against viral respiratory tract infection. In support of this, IgG with neutralizing potential has been shown in BAL samples from COVID-19 patients (Sterlin et al 2020). On the backdrop of observed antibody dependent enhancement (ADE) of disease for Spike based SARS-CoV vaccines in animals, scientific advice recommended avoiding a Th2 response, and to limit the induction of non-neutralizing antibodies (Lee et al. 2020, Lamberti et al. 2020). The VB2060 candidate is based on RBD and was here shown to mediate high levels of neutralizing antibodies and a dominant Th1 response, thereby theoretically reducing the risk of inducing vaccine associated enhancement of disease in humans (Halstead et al. 2020).

CD8^+^ T cells in VB2060-vaccinated mice were dominated by the presence of IFN-γ and TNF-α and indicate an effective cytotoxic T cell response specific for SARS-CoV-2 infected cells, whereas CD4^+^ T cells showed a predominant polyfunctional Th1 responses (defined by combined IFN-γ /TNF-α/IL-2 production). Recent studies have shown the importance of CD8^+^ T cell responses in controlling SARS-CoV-2 infection, with mild disease associated with CD8^+^ T cell responses in patients (Peng et al. 2020). Another study showed that high levels of SARS-CoV-2 responsive T cells were associated with protection from symptomatic SARS-CoV-2 infection among personnel at high risk of infection (i.e. healthcare providers, fire and police services) (Wyllie et al. 2020).

With the extensive range of vaccine technology platforms applied to SARS-CoV-2, it is vital to highlight the difference between the vaccine formats in relation to addressing the unmet needs outlined in the WHO TPP for vaccines against COVID-19 (WHO, 2020). The current study has shown that one dose of the VB10.COV2 candidate VB2060 induced rapid and high levels of neutralizing antibodies, CD8^+^ and Th1 CD4^+^ T cell responses that lasted for at least 3 months in mice, with a strong boost effect at day 89 indicating effective memory responses. Animal challenge studies are ongoing to inform further clinical development. The findings in this study, together with accrued safety data in humans on the similar vaccines from the same platform (Krauss et al. 2019, Hillemanns et al. 2020), the simplicity and scalability of plasmid DNA manufacturing, low cost of goods, preliminary data indicating long term storage at +2° to 8°C and simple administration, the VB2060 candidate is likely to be a promising future candidate to prevent COVID-19.

## METHODS

### Plasmid construction and testing of transient transfection in HEK293 cells

The VB10.COV2 constructs were designed as shown in Figure 1, and thereafter synthesized, cloned and produced by Genscript. The antigenic unit with either RBD or Spike was synthesized and cloned into a pUMVC4a VB10 master plasmid using SfiI-SfiI restriction enzyme sites. The resulting constructs encoded for homodimeric proteins with LD78β targeting units (Ruffini et al. 2010) and RBD/Spike as an antigenic unit, connected via a homodimerization unit consisting of exons from the hinge h1 and h4 and C_H_3 of human IgG3 (Figure 1). HEK293 cells (ATCC) were transiently transfected with VB10.COV2 DNA plasmids. Briefly, 2 ×10^5^ cells/well were plated in 24-well tissue culture plates with growth medium (DMEM, 10% FBS and 1% penicillin/streptomycin) and transfected with 1 μg VB10.COV2 DNA plasmids using Lipofectamine^®^ 2000 reagent under the conditions suggested by the manufacturer (Invitrogen, Thermo Fischer Scientific). The transfected cells were maintained for 3 days at 37°C with 5% CO_2_, and the cell supernatant was harvested. An ELISA was performed to verify the amount of VB10.COV2 protein produced by the HEK293 cells and secreted into the cell supernatant. Briefly, ELISA plates (MaxiSorp Nunc-immuno plates) were coated with 1 μg/ml of anti-C_H_3 (MCA878G, BioRad) in 1x PBS with 100 μl/well and plates were incubated overnight at 4°C. The microtiter wells were blocked by the addition of 200 μl/well 4% BSA in 1x PBS. 100 μl of cell supernatant from transfected HEK293 cells containing VB10.COV2 proteins were used. For primary detection antibody, either biotinylated anti-human MIP-1α (R&D Systems) or SARS-CoV-2/2019-nCoV Spike/RBD Antibody (1:1000) (Sino Biological) was used. Streptavidin-HRP (1:3000) or anti-rabbit IgG-HRP (1:5000) was added as secondary detection antibody. All incubations were carried out at 37°C for 1 hour (h), followed by 3x washing with PBS-Tween, except for the blocking step which was performed at RT for 1h followed by loading of supernatant after discardment of blocking buffer. 100 μl/well of TMB solution was added, and color development was stopped after 5-15 min adding 100 μl/well of 1 M HCl. The optical density at 450 nm was determined on an automated plate reader (Thermo Scientific Multiscan GO).

### Immunization of animals

6-week-old, female BALB/c mice were obtained from Janvier Labs (France). All animals were housed in the animal facility at the Radium Hospital (Oslo, Norway). All animal protocols were approved by the Norwegian Food Safety Authority (Oslo, Norway). Mice were given either one dose (day 0) or two doses (days 0 and 21) or three doses (day 0, 21 and 89), of DNA plasmid vaccine administrated to each *tibialis anterior* (TA) muscle by needle injection followed by AgilePulse *in vivo* electroporation (EP) (BTX, U.S.). Dose-response levels explored were 2.5 μg, 25 μg and 50 μg, or 3, 6, 12.5 and 25 μg of VB10.COV2 constructs. Blood sampling was performed on days 0, 7, 14, 20, 28, 42, 56, 70, 90 and 99, and spleens were collected on days 7, 14, 28, 90 and 99. Bronchoalveolar lavage (BAL) samples were collected on days 14, 21 and 28 by injection of one mL of sterile PBS into the lungs via the trachea, followed by three rounds of flushing.

### Anti-RBD IgG ELISA

The humoral immune response was evaluated in sera and bronchoalveolar lavages (BAL) collected at different time points (day 7, 14, 20, 28, 42, 56, 70, 90 or 99) after vaccination by an ELISA assay detecting total IgG specific for RBD from SARS-CoV2. ELISA plates (MaxiSorp Nunc-Immuno plates) were coated with 1 μg/ml recombinant RBD-His protein antigen (Cat. No. 40592-V08H, Sino Biological) in 1x D-PBS overnight at 4°C. Plates were blocked with 4% BSA in 1x D-PBS for 1 h at RT. Plates were then incubated with serial dilutions of sera or undiluted BAL samples for 2 h at 37°C. Plates were washed 3x and incubated with a 1:50,000 dilution of anti-mouse total IgG-HRP antibody (Southern Biotech) and incubated for 1h at 37°C. After final wash, plates were developed using TMB substrate (Merck, cat. CL07-1000). Plates were read at 450 nm within 30 min using a Multiscan GO (Thermo Fischer Scientific). Binding antibody endpoint titers were calculated as the reciprocal of the highest dilution resulting in a signal above the cutoff. For BAL, responses were reported as OD_450_ values.

### SARS-COV-2 live neutralization assay

Live virus microneutralization assays (MNA) were performed at Public Health England (Porton Down, UK) as described (Folegatti et al. 2020). Neutralising virus titres were measured in heat-inactivated (56°C for 30 min) serum samples. Diluted SARS-CoV-2 (Australia/VIC01/20202) (Caly et al. 2020) was mixed 50:50 in 1% FCS/MEM with doubling serum dilutions in a 96-well V-bottomed plate and incubated at 37°C in a humidified box for 1 hour. The virus/serum mixtures were then transferred to washed Vero E6 (ECACC 85020206) cell monolayers in 96-well flat-bottomed plates, allowed to adsorb at 37°C for a further hour, before removal of the virus inoculum and replacement with overlay (1% w/v CMC in complete media). The box was resealed and incubated for 24 hours prior to fixing with 8% (w/v) formaldehyde solution in PBS. Microplaques were detected using a SARS-CoV-2 antibody specific for the SARS-CoV-2 RBD Spike protein and a rabbit HRP conjugate, infected foci were detected using TrueBlueTM substrate. Stained microplaques were counted using ImmunoSpot^®^ S6 Ultra-V Analyzer and resulting counts analysed in SoftMax Pro v7.0 software. International Standard 20/130 (human anti-SARS-CoV-2 antibody from human convalescent plasma, NIBSC, UK) was used for comparison.

### IFN-γ ELISpot assay

Splenocytes from vaccinated mice were analyzed in IFN-γ ELISpot assay detecting RBD-/Spike-specific T cell responses. Briefly, the animals were sacrificed at day 7, 14, 28, 90 or 99, and the spleens were harvested aseptically. The spleens were homogenized, single-cell suspensions were incubated with 1 × ACK buffer to remove erythrocytes, washed and re-suspended to a cell concentration of 6 × 10^6^ cells. CD4 or CD8 T-cell populations were depleted from the total splenocyte population using the Dynabead (catalog no. 11447D or 11445D, Thermo Fischer Scientific) magnetic bead system according to the manufacturer’s recommended procedures. Cells were then re-suspended at 6 x 10^6^ cells/ml for the ELISpot assay. Depletion was confirmed by flow cytometry. The cells were plated in triplicates (6 x 10^5^ cells/well) and stimulated with 2 μg/ml of RBD/Spike peptide pools (RBD: 6 pools consisting of 10-11 x of 15 mers overlapping with 12 aa and Spike: 24 pools consisting of 12 x 15 mers overlapping with 11 aa) or individual peptides for 24h. Cells without peptide stimulation was used as negative controls. The stimulated splenocytes were analyzed for IFN-γ responses using the mouse IFN-γ ELISpot Plus kit (Mabtech AB, Sweden). Spot-forming cells (SFU) were measured in a IRIS™ ELISpot reader using the APEX™ software from Mabtech AB, Sweden. Results are shown as the mean number of IFN-γ + spots/10^6^ splenocytes with subtracted background.

### Cell stimulation and staining for flow cytometry

The animals were sacrificed 28 days post the first dose and spleens were removed aseptically. The spleens were mashed to obtain single-cell suspensions, and 1x ACK buffer was used to remove erythrocytes. The splenocytes were then washed, plated (2.0 x 10^6^ cells/well in 24 well plates) and stimulated for 16 h with 6 μg/ml of RBD peptide pools. For detection of cytokines with flow cytometry, 1x monensin and 1x brefeldin were added to the wells 1h post incubation start. Following the stimulation with RBD peptide pools, the cells were harvested, and centrifuged twice with PBS to wash away the medium. The cells were incubated with fixable viability dye (eFluor780) in the dark for 10 min at RT. Cells were further stained with the extracellular antibodies (anti-CD3, anti-CD4, anti-CD8 and γδTCR), fixed and permeabilized, and stained for detection of cytokines (anti-TNFα, anti-IFN-γ, anti-IL-4, anti-IL-17 antibodies) and a transcription factor (anti-FoxP3 antibody). Detailed description of antibodies used for flow cytometry are shown in Supplemental Table S3. The stained cells were run in BD FACSymphony A5 (BD Biosciences, U.S.) and analyzed using FlowJo software.

### Multiparameter flow cytometry analysis

The RBD-stimulated mice splenocyte T cells were defined through the exclusion of dead cells, doublets and CD3^-^ non-T cells (Figure S4A-D). CD3^+^ T cells were then analyzed for the presence of γδTCR T cells and these cells were further removed from the analysis (Figure S4E). The remaining T cells were then examined for CD4^+^ and CD8^+^ markers, and the majority expressed either CD4^+^ or CD8^+^ T cell populations (Figure S4F). Both populations were examined for individual expression of IFN-γ, TNF-α, IL-4, IL-17 or FoxP3 and gates were set to define positive cells. These positive cells were further analyzed using Boolean gating algorithm in FlowJo software. The algorithm calculated all possible combinations of cytokines produced by each cell, thus allowing analysis of multifunctional T cells on a single-cell level.

### Cytokine release testing

Cell culture supernatant from splenocytes stimulated with RBD peptides for 16 h was harvested and analyzed for cytokine presence. In short, 50 μl of the cell culture supernatant was used as described in the supplier’s protocol for Essential Th1/Th2 Cytokine 6-Plex Mouse ProcartaPlex™ Immunoassay (Thermo Fisher Scientific). Presence of IFN-γ, TNF-α and IL-12p70 in the supernatant defined Th1 response. The Th2 response was defined through the production of IL-4 and IL-5.

### Statistical analysis

Statistical analyzes of antibody responses in sera to compare groups were performed by two-tailed Mann-Whitney test was performed (GraphPad Software). A value of p< 0.05 was considered significant.

### Data availability

The data that support the findings of this study are available from the corresponding author, GN, upon reasonable request.

## ACKNOWLEDGEMENTS

Audun Bersaas and Renate Skarshaug are thanked for excellent work in supporting and performing laboratory analyses, and the Vaccibody team for project support. We also want to thank the staff at the animal research facility at the Oslo Radium hospital. We also thank the staff at Oslo Radium Hospital Flow Cytometry Core Facility for the technical support. We thank PHE, Porton Down, U.K., for neutralization assay testing performed as a service in collaboration with Nexelis, Canada.

## AUTHOR CONTRIBUTIONS

ABF, MS, GN, ES, LMS, BS conceptualized experiments. ES, LMS, BS, SC, RS, LB, KK, AB and EM performed the experiments. GN, ES and ABF wrote the manuscript. All authors supported the review of the manuscript.

## ADDITIONAL INFORMATION

Supplementary information is available for this paper (Figures and Tables denoted “S”). The research is funded by Vaccibody. Correspondence and requests for materials should be addressed to

## Supplemental Materials

**Supplemental figure S1.**
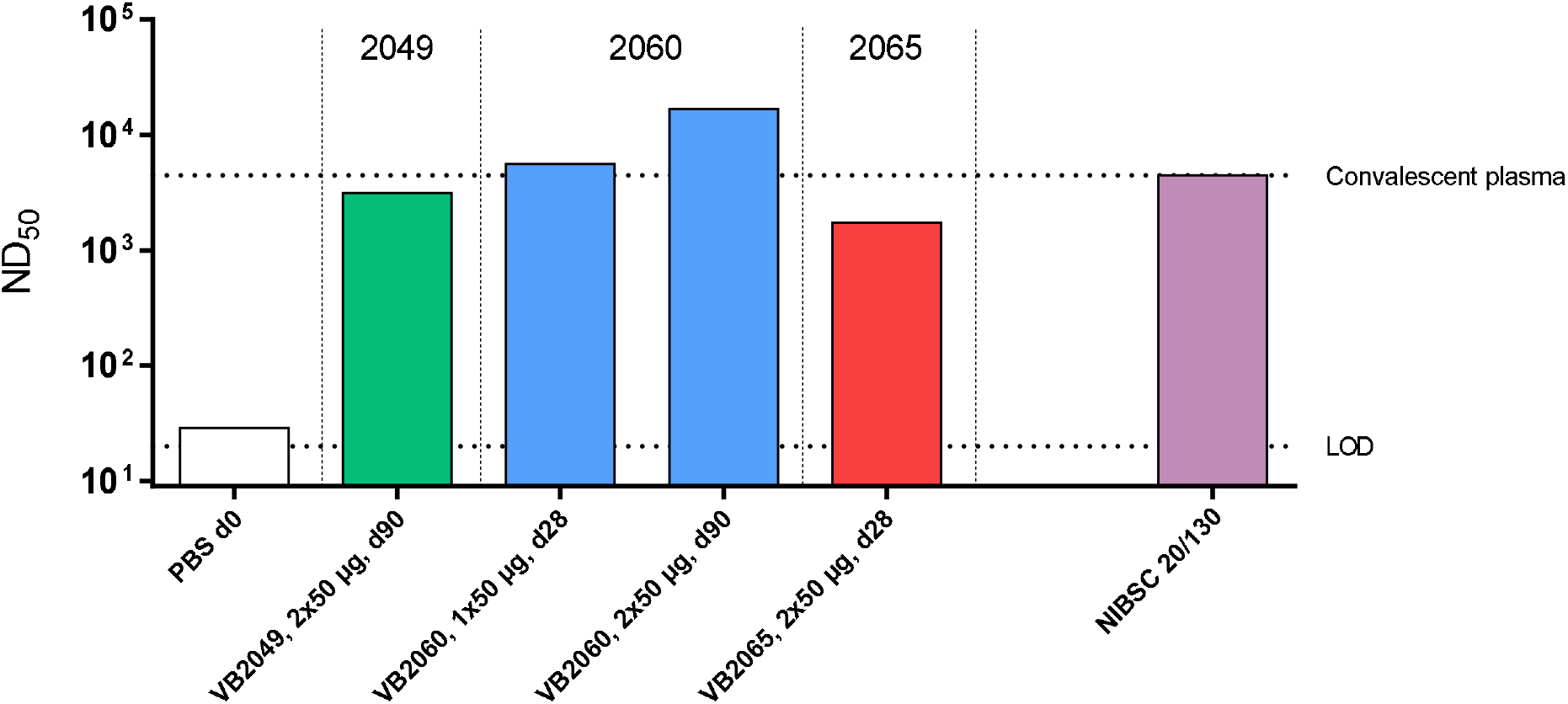
Neutralizing antibody responses in sera after immunization with VB2049, VB2060 and VB2065. Bars represent endpoint titers for pooled sera assessed in live virus neutralization against homotypic SARS-CoV-2 live virus strain Australia/VIC01/2020. Lower dashed line indicate the limit of detection (LOD) of the assay, and the upper dashed line the titer value for the convalescent serum reference NIBSC 20/130 (ND50 endpoint titer 4443). Due to limited serum volume available, only pools from selected timepoints could be tested.

**Supplemental figure S2.**
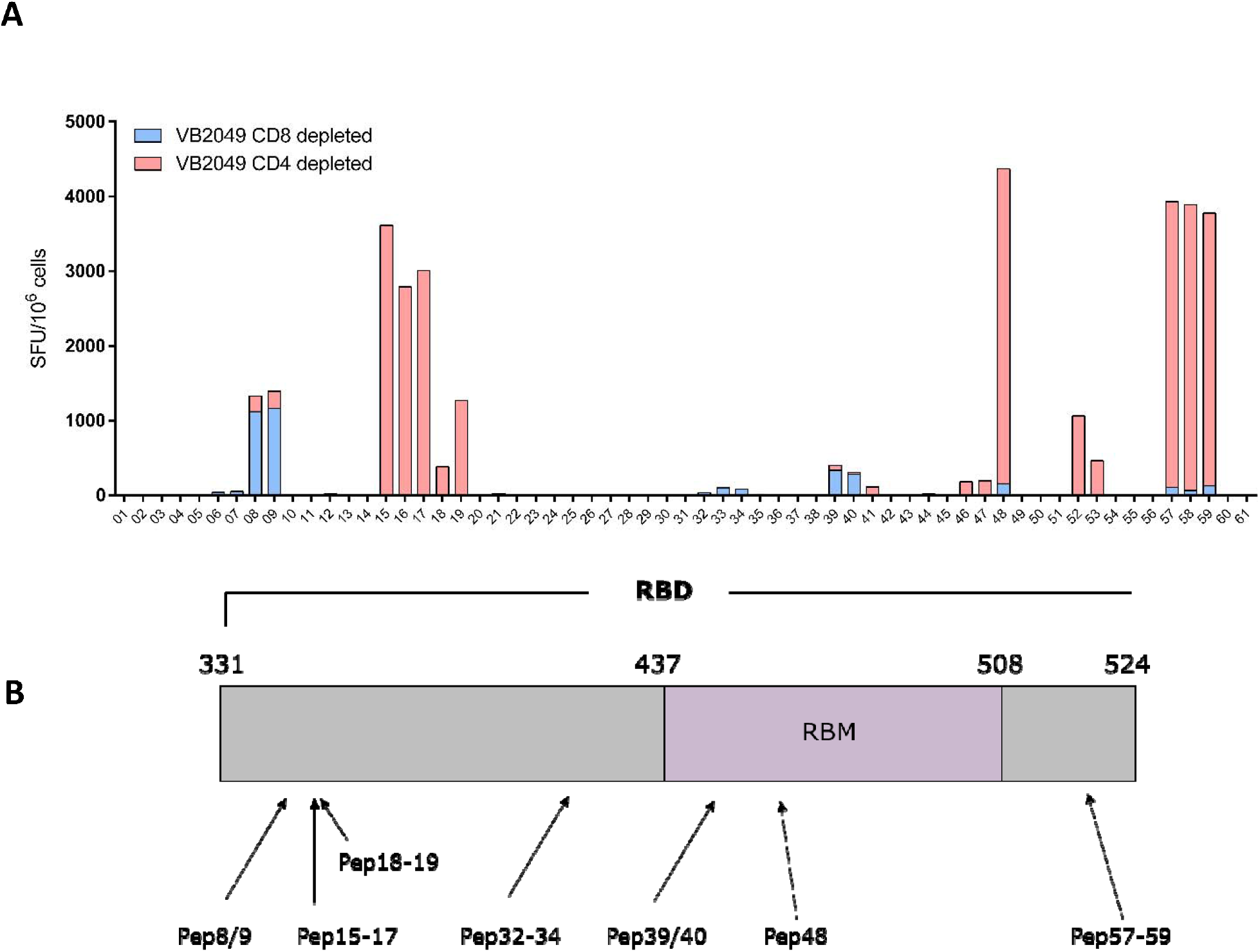
Identification of CD4^+^ and CD8^+^ RBD-specific T cell epitopes following i.m. vaccination with 25 μg VB2049 at day 0 and day 21 in mice (*n* = 5). **a** CD4^+^ and CD8^+^ T cell populations were stimulated for 24 h with 61 individual RBD peptides (15-mer peptides overlapping by 12 aa from SARS-CoV2 RBD domain) and the number of IFN-γ positive spots/1×10^6^ splenocytes were detected in an ELISpot assay 1 week post boost vaccination. **b** Map of the SARS-CoV2 RBD domain and identification of immunodominant peptides in mice. RBM; receptor-binding motif.

**Supplemental figure S3.**
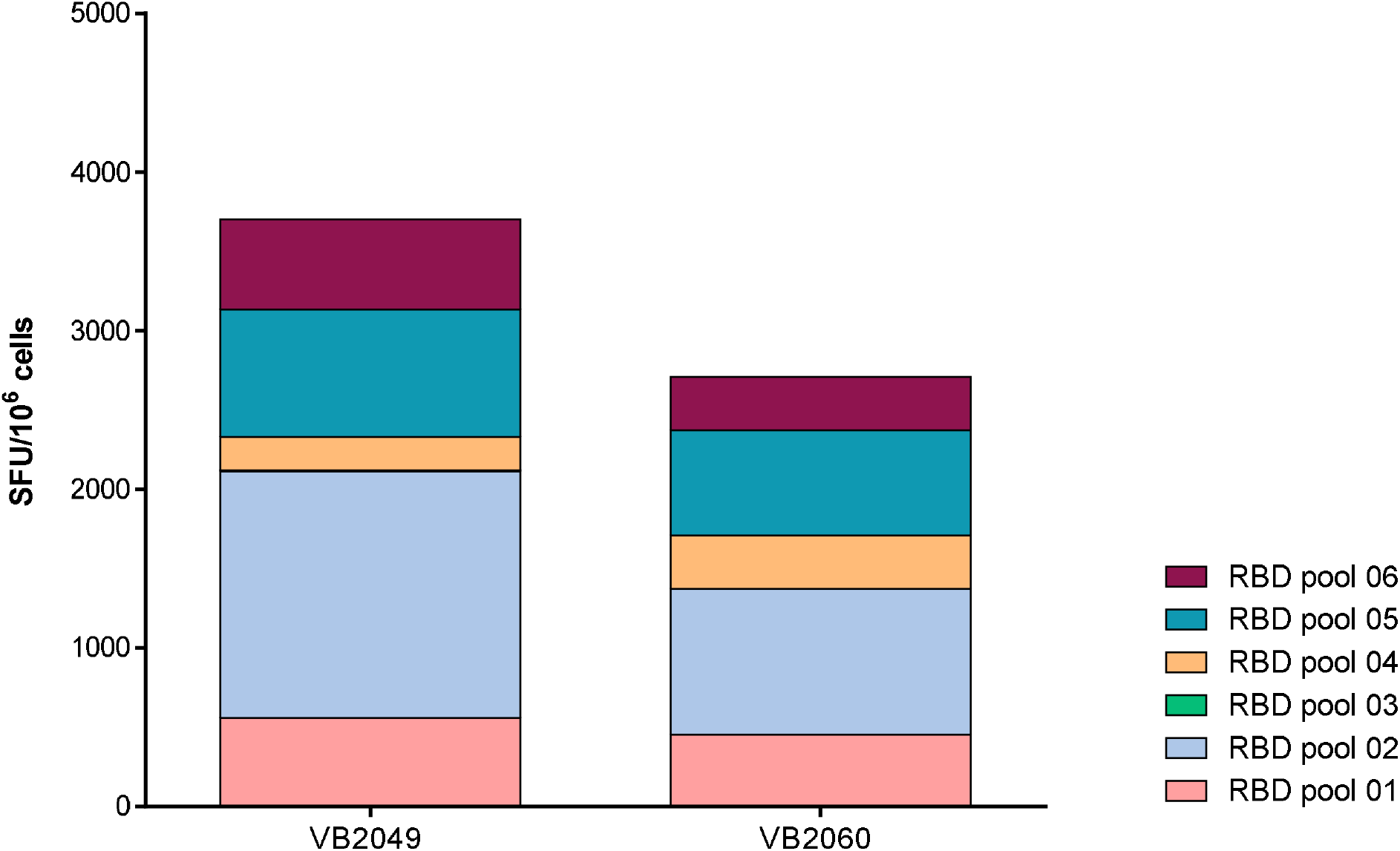
Comparative T cell responses induced by VB10.COV2 DNA vaccines VB2049 and VB2060. Two doses of 2.5 μg vaccine administered i.m. at days 0 and 21, and splenocytes were harvested at day 28 post first immunization. The total number of IFN-γ positive spots/1×10^6^ splenocytes after restimulation with overlapping RBD peptide pools were determined by IFN-γ ELISpot.

**Supplemental figure S4.**
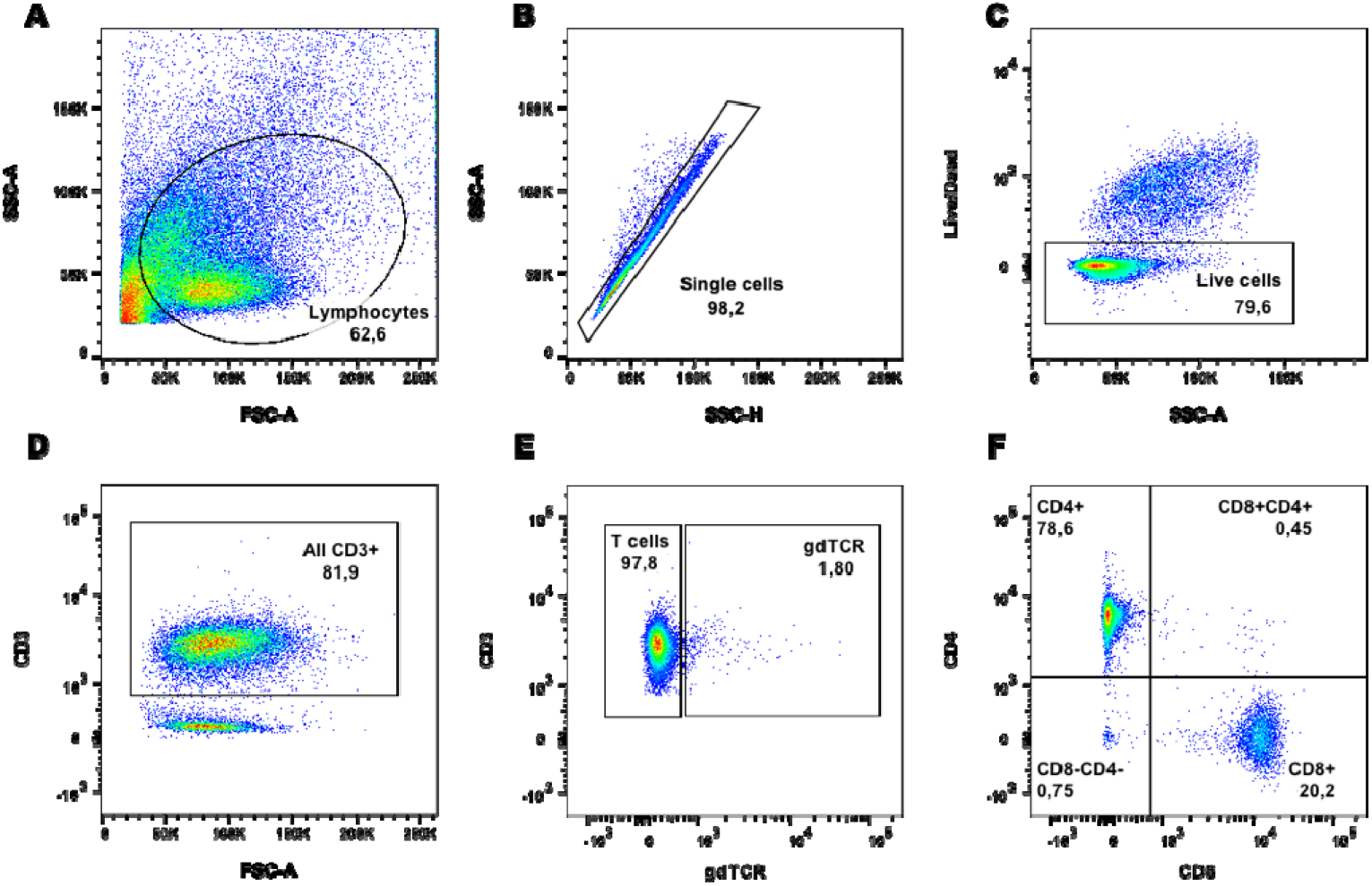
Gating strategy for the identification of T cells. **a** All cells were examined using side scatter (SSC) and forward scatter (FSC) parameters. Lymphocyte gate was set based on the relative size (FSC) of the cells. **b** Lymphocytes were analyzed for presence of doublets, and a gate was set to include only single cells in further analysis. **c** Dead cells were identified using viability dye and a gate was set to include live cells in further analysis. **d** In the population of live cells, all CD3^+^ cells were gated for future analysis. **e** T cells were defined as CD3+ and γδTCR T cells were excluded from the analysis. **f** All T cells were analyzed for expression of CD4 and CD8 markers.

**Supplemental table S1.**
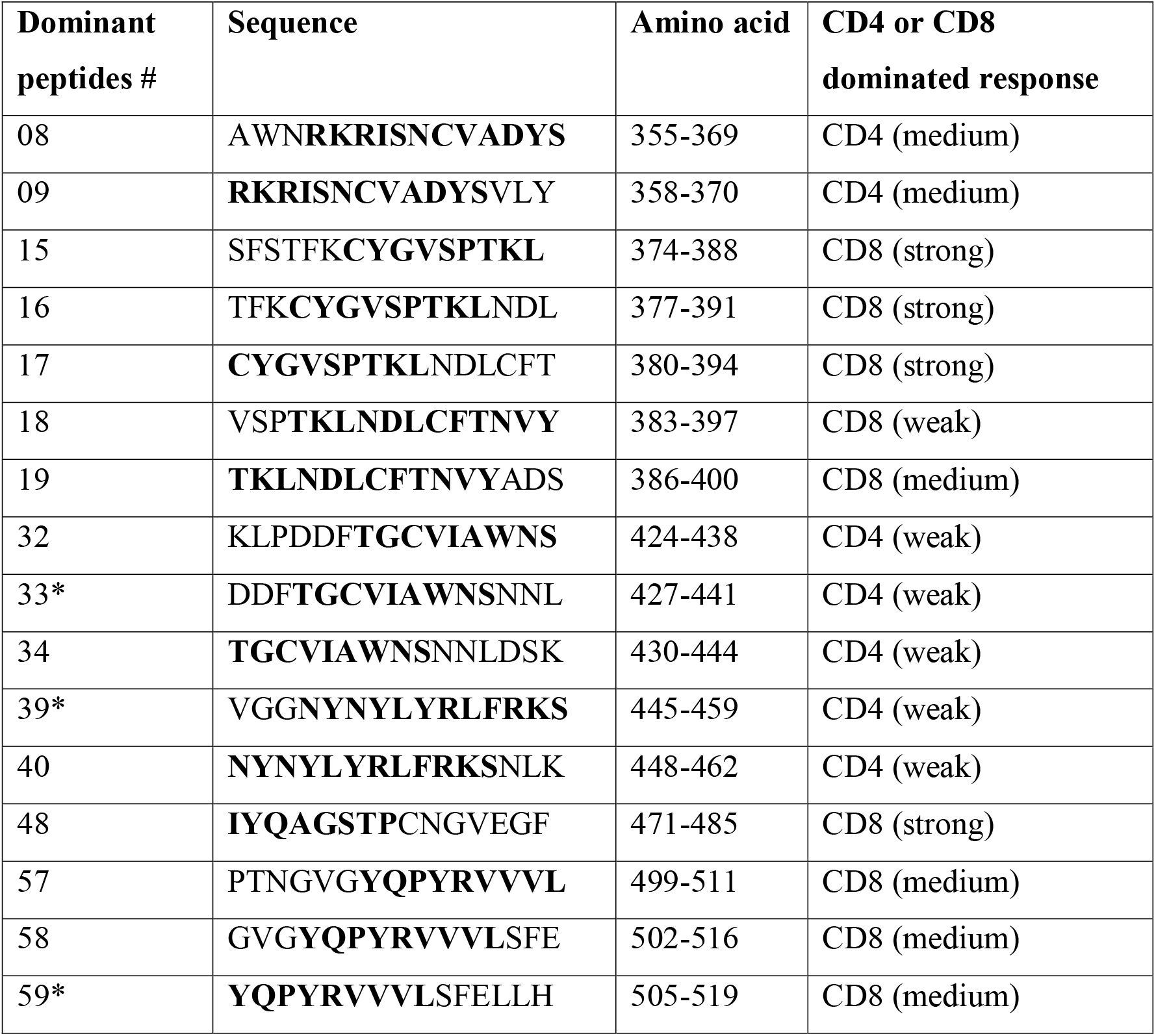
Individual peptides tested for induction of CD4+ and CD8+ RBD specific immune responses and T cell epitope mapping after i.m. vaccination with 25 μg of VB2049 in mice. Sequences that were immunodominant in mice (i.e. strong CD4^+^ or CD8^+^ responses) are shown in shaded columns. *; epitopes verified in a separate study mapping T cell epitopes induced by a Spike based DNA vaccine candidate in mice (Smith et al. 2020).

**Supplemental table S2.** Specificity of peptides eliciting significant T cell responses by VB2065 (Spike).

| **Spike pool** | **Peptides** | **aa** | **Position in Spike** |
| --- | --- | --- | --- |
| Pool 06 | 62-73 | 244-297 | NTD (N-terminal domain) |
| Pool 11 | 123-134 | 488-541 | RBD (receptor binding domain) |
| Pool 20 | 231-238,240-242,244,245 | 920-986 | HR1 (heptapeptide repeat sequence 1) |
| Pool 22 | 257-269 | 1024-1081 |  |
| Pool 23 | 270-276,278,280-283 | 1076-1137 |  |

**Supplemental table S3.** Description of antibodies used for flow cytometry.

| Target | Fluorochrome | Vendor | Catalogue No. |
| --- | --- | --- | --- |
| CD3 | Brilliant ultraviolet 395 | BD Bioscience | 740268 |
| CD4 | Brilliant violet 785 | Biolegend | 100453 |
| CD8 | PE/Cyanine 7 | Biolegend | 100721 |
| $\gamma\delta$TCR | PerCP/Cyanine5.5 | Biolegend | 118117 |
| TNF-α | Brilliant violet 605 | Biolegend | 506329 |
| IFN-γ | APC | Biolegend | 505809 |
| IL-4 | PE | Biolegend | 504103 |
| IL-2 | Brilliant violet 421 | Biolegend | 503825 |
| IL-17 | Alexa Fluor 488 | Biolegend | 506909 |
| FoxP3 | Alexa Fluor 700 | Biolegend | 126421 |

